# Estrogen induces mammary ductal dysplasia

**DOI:** 10.1101/673525

**Authors:** Junji Itou, Rei Takahashi, Hiroyuki Sasanuma, Masataka Tsuda, Suguru Morimoto, Yoshiaki Matsumoto, Tomoko Ishii, Fumiaki Sato, Shunichi Takeda, Masakazu Toi

**Affiliations:** Laboratory of Molecular Life Science, Institute for Biomedical Research and Innovation, Foundation for Biomedical Research and Innovation at Kobe (FBRI), 2-2 Minatojima-Minamimachi, Chuo-ku, Kobe 650-0047, Japan; Department of Breast Surgery, Graduate School of Medicine, Kyoto University, 54 Shogoin-Kawahara-cho, Sakyo-ku, Kyoto 606-8507, Japan; Graduate School of Pharmaceutical Sciences and Faculty of Pharmaceutical Sciences, Doshisha Women’s College of Liberal Arts, 97-1 Kodo, Kyotanabe 610-0395, Japan; Department of Radiation Genetics, Graduate School of Medicine, Kyoto University, Yoshida-Konoe-cho, Kyoto 606-8501, Japan; Program of Mathematical and Life Science, Graduate School of Integrated Sciences for Life, Hiroshima University, 1-3-1 Kagamiyama, Higashi-Hiroshima 739-8526, Japan; Department of Breast Surgery, Kansai Electric Power Hospital & Kansai Electric Power Medical Research Institute, 2-1-7 Fukushima, Fukushima-ku, Osaka 553-0003, Japan

**Keywords:** Breast cancer, Dysplasia, Estrogen, Isoflavones

## Abstract

Mammary ductal dysplasia is a phenotype observed in precancerous lesions and early-stage breast cancer. However, the mechanism of dysplasia formation remains elusive. Here we show, by establishing a novel dysplasia model system, that estrogen, a female hormone, has the potential to cause mammary ductal dysplasia. We injected estradiol (E2), the most active form of estrogen, daily into scid mice with a defect in nonhomologous end joining repair and observed dysplasia formation with cell proliferation at day 30. Moreover, we found that isoflavones inhibited E2-induced dysplasia formation. Our dysplasia model system provides insight into the mechanistic understanding of breast tumorigenesis, and the development of breast cancer prevention.

## Introduction

During breast tumorigenesis, mammary ductal dysplasia is observed in precancerous lesions and early-stage breast cancers. Mammary ductal dysplasia exhibits a loss of the biphasic mammary epithelial and myoepithelial pattern, an abnormal nucleus, epithelial cell expansion, a disruption in the myoepithelial cell layer and/or mammary epithelial cell invasion to fibrous stroma, whereas a normal mammary duct maintains cell polarization, the biphasic pattern and the smooth luminal surface of the mammary epithelial cell layer. Given that dysplasia can progress to malignant neoplasms (1–3), elusicating the mechanism of dysplasia formation will contribute to the prevention of breast tumorigenesis.

Previous studies have established various breast cancer mouse models by genetic engineering. For example, the mammary gland-specific expression of *c-neu* (*Her2*/*ErbB2*), *polyoma middle T* antigen and *Wnt-1* causes breast cancer (4–6). Mice with a mutation in *Tp53*, a tumor suppressor gene, also developed breast cancer (7). Although these models have provided knowledge about breast cancer, particularly at an advanced stage, they have not been primarily used for studies on mammary ductal dysplasia observed in precancerous lesions and early-stage breast cancers. Previous dysplasia studies in mice used radiation in *Atm* heterozygous mice (8), and overexpression of constitutively activated SMO (9) and of the oncogene NSD3 (10). Because studies using genetically engineered mice have been designed to examine phenotypes specifically caused by the functions of their target genes, these models are not suitable to elucidate the mechanism by which mammary ductal dysplasia is naturally formed. To understand such a mechanism of dysplasia formation, a mouse model that form dysplasia by physiological factor(s) is required.

Estrogen, a female hormone, promotes the development of the normal mammary dust and the proliferation of breast cancer (reviews, (11–14)). There are a number of studies on the function of estrogen in breast cancer (reviews, (11, 13–16)). Although estrogen is thought to be involved in breast tumorigenesis in epidemiologic studies (17), and combination of radiation and E2 treatment transformed normal mammary cells *in vitro* (18), there is no experimental evidence showing that estrogen receptor α (ERα)-mediated estrogen signaling induces mammary dysplasia from normal mammary epithelial cells *in vivo*.

Estrogen regulates gene expression via the activation of ERα. In mammary glands, ERα is expressed mainly in mammary epithelial cells. Although there are various functional models of ERα action, in the classical mechanism of ERα, estrogen-bound ERα forms a dimer, localizes in the nucleus, and binds to its DNA binding site to regulate gene expression (reviews, (12, 15, 19–21)). Thus far, various ERα-regulated genes have been identified (13, 14, 16); these include *growth regulating estrogen receptor binding 1* (*GREB1*), *trefoil factor 1* (*TFF1*, *pS2*) (22, 23). During ERα-mediated gene regulation, a DNA double-strand break is made at the promoter of a target gene to promote its expression (24, 25). Whether ERα-mediated gene regulation with a DNA double-strand break is involved in dysplasia formation remains unclear. In this study, we successfully established a novel dysplasia model system and investigate the mechanism of dysplasia formation.

## Results

### Enhancement of ERα-mediated estrogen signaling causes mammary ductal dysplasia

To investigate E2-induced DNA damage *in vitro*, We treated an ERα-positive breast cancer cell line, MCF-7, with or without estradiol (E2), the most active form of estrogen, and investigated DNA double-strand breaks (Fig. S1A). We counted the number of the signals of phosphorylated-histone H2AX (gamma-H2AX, gH2AX), a marker of DNA double-strand breaks. The number of gH2AX signals was increased by E2 treatment and reduced by cotreatment with fulvestrant, an estrogen receptor inhibitor (Fig. S1A), indicating that E2 treatment causes DNA double-strand breaks via its receptor. An inhibitor of DNA-dependent protein kinase (DNA-PK), NU-7441, inhibits nonhomologous end joining repair, one of the repair mechanisms of DNA double-strand breaks (26). NU-7441 treatment did not change the number of gH2AX signals after 2 h treatment (Fig. S1A), indicating that the loss of DNA-PK function does not change E2-generated DNA damage in our conditions.

To determine whether estrogen-induced DNA double-strand breaks are involved in the regulation of the downstream genes of estrogen signaling, we reduced the capacity of nonhomologous end joining by knocking down *PRKDC*, which encodes the catalytic subunit of DNA-PK (27). Cells were washed with medium in the absence of E2 after E2 treatment to analyze repair capacity. No increase in the number of gH2AX signals was observed in control cells (Fig. 1A), indicating that cells repaired E2-induced DNA double-strand breaks during 2 h of washing under our conditions. However, the gH2AX signal was retained after washing in *PRKDC* knockdown cells (Fig. 1A, S1B), suggesting that the loss of DNA-PK function delays the repair of E2-induced DNA double-strand breaks. Longer-retaining DNA breaks are believed to be involved in transcriptional regulation, at least in specific cases (28). To investigate the expression of ERα downstream genes under our conditions, mRNA levels were quantified (Fig. 1B). The expression of *GREB1* and *TFF1* tended to increase following *PRKDC* knockdown, although there was no significant difference. These results suggest that a reduction in the repair capacity for DNA double-strand breaks enhances the expression of some ERα-regulated genes in response to E2 stimulation.

**Fig. 1.**
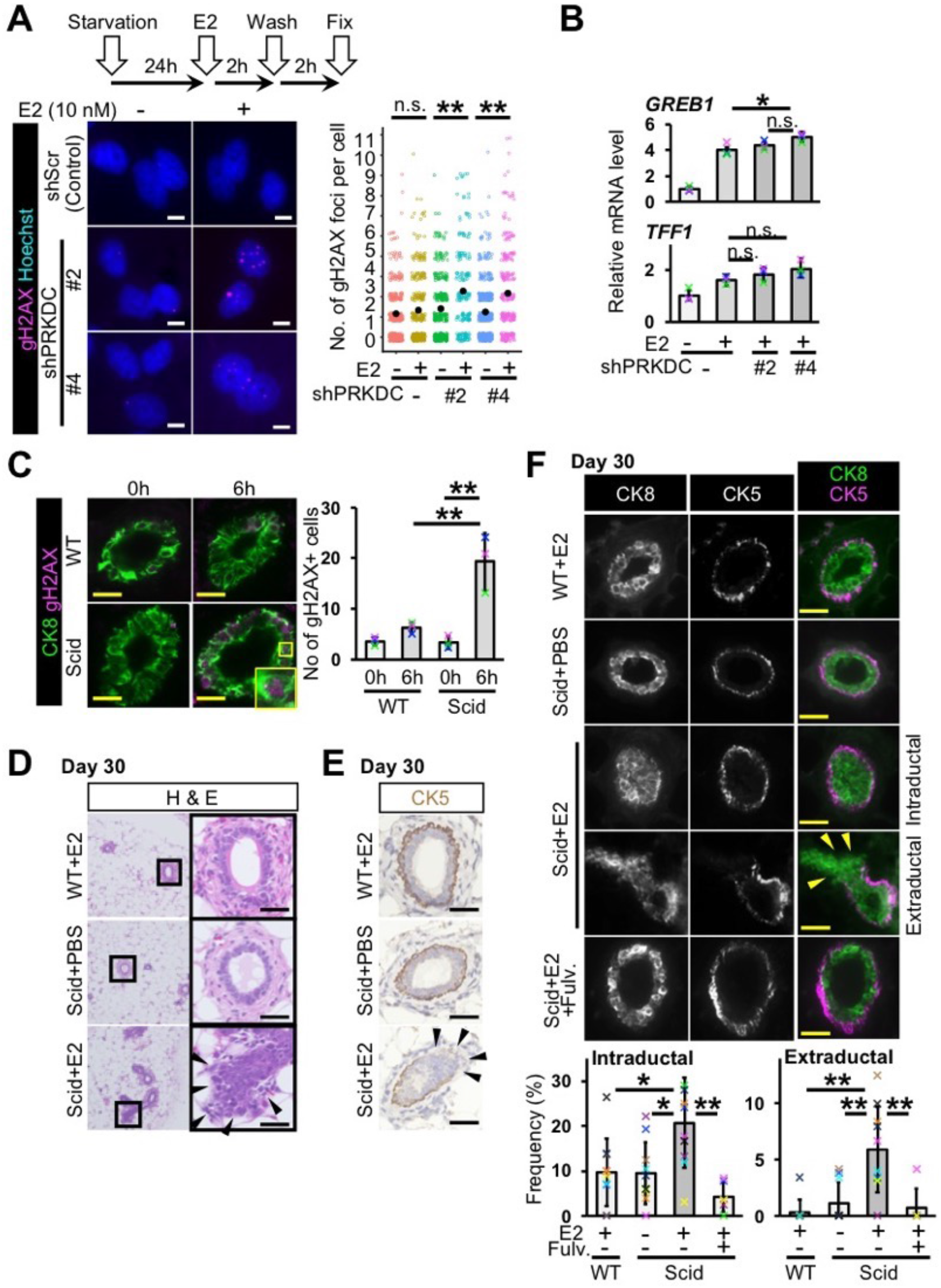
Estrogen administration induces mammary ductal dysplasia in scid mice. A, DNA double-strand breaks were detected in MCF-7 cells. *PRKDC* was knocked-down. Gamma-H2AX was immunostained. Numbers of gH2AX foci per cell were graphed (jitter plot). Black dots indicate mean values. Data were obtained from 2 or 3 independent experiments (total 200~520 cells in each group, Mann-Whitney U test). B, Messenger RNA levels of *GREB1* and *TFF1* were quantified (*n*=3 experiments, Tukey’s test). Cells were treated with or without E2 for 6 h. C, Gamma-H2AX-positive mammary epithelial cells were detected and quantified at 6 hours after E2 administration (*n*=3 mice, Tukey’s test). D, Typical images of H&E staining are shown. Daily injection of E2 was performed for 30 days. E, Typical immunostaining images of CK5 are shown. F, Fluorescent images of CK8 and CK5 staining are shown. Mammary ducts with intraductal and extraductal expansion were quantified (*n*=10 mice in WT+E2, Scid+PBS and Scid+E2 groups, *n*=6 mice in Scid+E2+Fulv. group, Tukey’s test and Mann-Whitney U test). Fulv.: fulvestrant. Scale bars indicate 10 μm (A) and 30 μm (C-F). n.s.: not significant, *: *P*<0.05, **:*P*<0.01. Error bars represent standard deviation. Arrowheads indicate mammary epithelial cells in extraductal region (D-F).

We injected E2 (6 μg/mouse) intraperitoneally into scid mice with a genetic defect in DNA-PKcs function (29, 30). The serum concentration of E2 was transiently increased and reduced to the background level after 6 h in both wild-type and scid mice (Fig. S1C). The results of gH2AX staining showed an increased number of gH2AX-positive mammary epithelial cells (Fig. 1C). As expected, scid mice showed a larger number of gH2AX-positive cells than wild-type mice.

To investigate whether long term-E2 administration could cause an abnormality in the mammary gland, we performed consecutive daily injections of E2 for 30 days. No significant change in body weights was observed after 30 days of injection (Fig. S2A). Hematoxylin and eosin (H&E) staining showed normal mammary ducts in both wild-type and scid mice at day 7 (Fig. S2B). At day 30, whereas E2-injected wild-type (WT+E2) and phosphate-buffered saline (PBS)-injected scid (scid+PBS) mice exhibited normal mammary ducts (Fig. 1D), dysplasia formation (i.e., increased cell number, loss of biphasic mammary epithelial and myoepithelial patterns, and epithelial cell expansion to outside of a duct) was observed in E2-injected scid mice (scid+E2) (Fig. 1D,S2C). Because human breast cancer shows a disruption in the myoepithelial layer (3), we immunostained cells for the myoepithelial cell markers cytokeratin 5 (CK5) (Fig. 1E) and p63 (Fig. S2D). We observed a disruption in the myoepithelial layers in mammary ducts with dysplasia. To determine whether mammary epithelial cells of the ducts with dysplasia invaded to the outside of myoepithelial layer, co-immunostaining of CK5 and a mammary epithelial cell marker CK8 was performed (Fig. 1F). The results revealed mammary ducts with increased mammary epithelial cell numbers and loss of the biphasic pattern (Fig. 1F intraductal) and mammary ducts with epithelial cells in the extraductal region with a disruption in the CK5 cell layer (Fig. 1F extraductal) in scid+E2. Since terminal end buds are epithelial cell-rich regions that can be identified as a duct with no fibrous stroma region (31) and since it is difficult to distinguish dysplasia from normal terminal end bud, these were excluded in the analyses. We analyzed the frequency of dysplasia formation and observed significant increases in both intraductal and extraductal expansion (Fig. 1F graph). A duct with both intraductal and extraductal expansion was counted as a duct with extraductal expansion. These results suggest that repetitive E2 administration induces mammary ductal dysplasia in mice that are susceptible to estrogen, such as scid mice. Coadministration with the estrogen receptor inhibitor (scid+E2+Fulv.) showed no increase in the frequency of dysplasia, suggesting that the suppression of estrogen receptor function may prevent dysplasia formation.

We coadministered NU-7441, a DNA-PK inhibitor, and E2 to two wild-type strains, C.B17/Icr and C57BL/6J. C.B17/Icr is the parental strain of scid mice. C57BL/6J is a commonly used strain. At day 30, we observed an increase in dysplasia formation in NU-7441+E2 mice compared to NU-7441+PBS mice in both strains (Fig. S3). This result indicates that E2 administration can induce mammary ductal dysplasia not only in scid mice but also in various mouse backgrounds.

### Progesterone inhibits E2-induced dysplasia formation

Progesterone (PG), a female hormone, alters ERα function in malignant breast cancer through progesterone receptor (PGR), and the administration of PG reduced E2-dependent tumor growth in mouse xenograft experiments with MCF-7 cells (32). To determine whether PG-activated PGR inhibits E2-induced DNA double-strand breaks in MCF-7 cells, we performed combination treatment of E2 and PG with or without *PGR* knockdown (Fig. 2A, B). The number of gH2AX signals was increased by E2 treatment but not in the combination of E2 and PG (Fig. 2B). In *PGR* knocked-down cells, the combination treatment of E2 and PG resulted in increased number of gH2AX signals, similar to E2 treatment, suggesting that PGR prevents E2-induced DNA double-strand breaks.

**Fig. 2.**
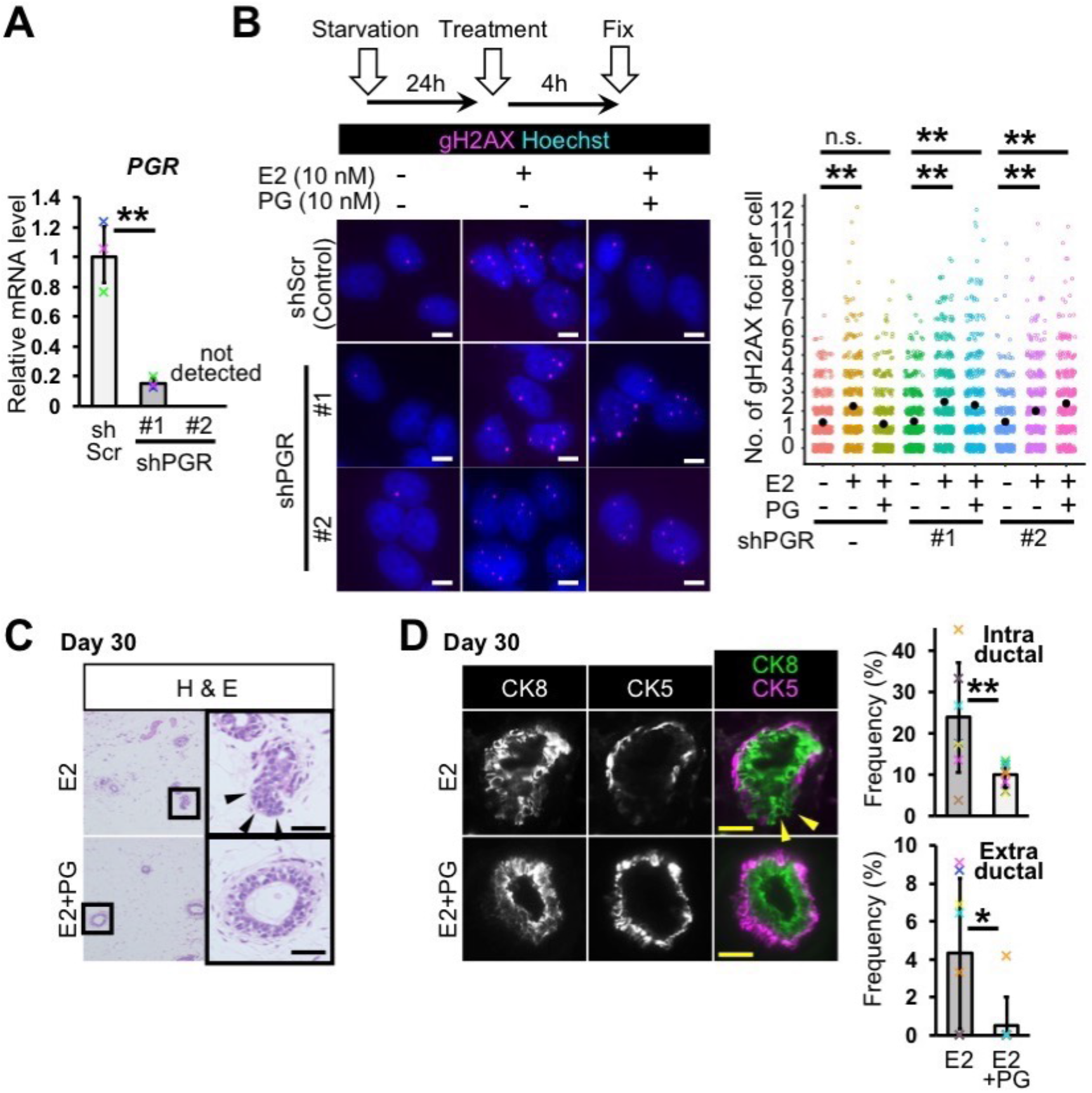
Progesterone inhibits estrogen-induced mammary ductal dysplasia. A, Progesterone receptor gene was knocked-down (*n*=3 experiments, student’s *t*-test to shScr control). B, DNA double-strand breaks were detected by gH2AX immunostaining. Numbers of gH2AX foci per cell were analyzed (jitter plot) (*n*=3 independent experiments, total 380~520 cells in each group, Mann-Whitney U test). C, Typical images of H&E staining are shown. D, Fluorescent images of CK8 and CK5 staining are shown. Mammary ducts with intraductal and extraductal expansion were quantified (*n*=8 mice, Mann-Whitney U test). Scale bars indicate 10 μm (B) and 30 μm (C,D). n.s.: not significant, *: *P*<0.05, **:*P*<0.01. Error bars represent standard deviation. Arrowheads indicate mammary epithelial cells in extraductal region (C,D).

The function of PG in dysplasia formation *in vivo* remain elusive. To this end, we coinjected E2 and PG into our dysplasia model system. In scid mice, the administration of E2 alone caused dysplasia formation, and the coadministration of E2 and PG prevented dysplasia formation (Fig. 2C). Double immunostaining of CK5 and CK8 showed that PG administration prevented extraductal expansion (Fig. 2D). The quantification of intraductal and extraductal expansion showed that PG reduced the frequencies of these abnormalities (Fig. 2D). These results suggest that PG has a protective effect for E2-induced DNA damage and dysplasia formation.

### Estrogen promotes mammary epithelial cell proliferation

Because the number of CK8-positive cells was likely increased in scid+E2 mice (Fig. 1F), we investigated cell proliferation by immunostaining for PCNA, an S-phase marker, and Ki-67, a proliferation marker, at day 30 (Fig. 3A, B). The results of immunostaining showed that the ratio of proliferating mammary epithelial cells was increased in scid+E2 mice, and this increase was inhibited by the ERα inhibitor (scid+E2+Fulv.). An increased number of proliferating cells was also observed at day 7 (Fig. 3C). To investigate organ level changes, we visualized mammary ducts by Carmine Alum-staining (Fig. 3D). The mammary glands of scid+E2 mice were more densely distributed than those of the control mice and exhibited more small small branches. We quantified the number of branches and found that it was increased in scid+E2. These results suggest that E2 administration induces cell proliferation in the mammary glands of scid mouse.

**Fig. 3.**
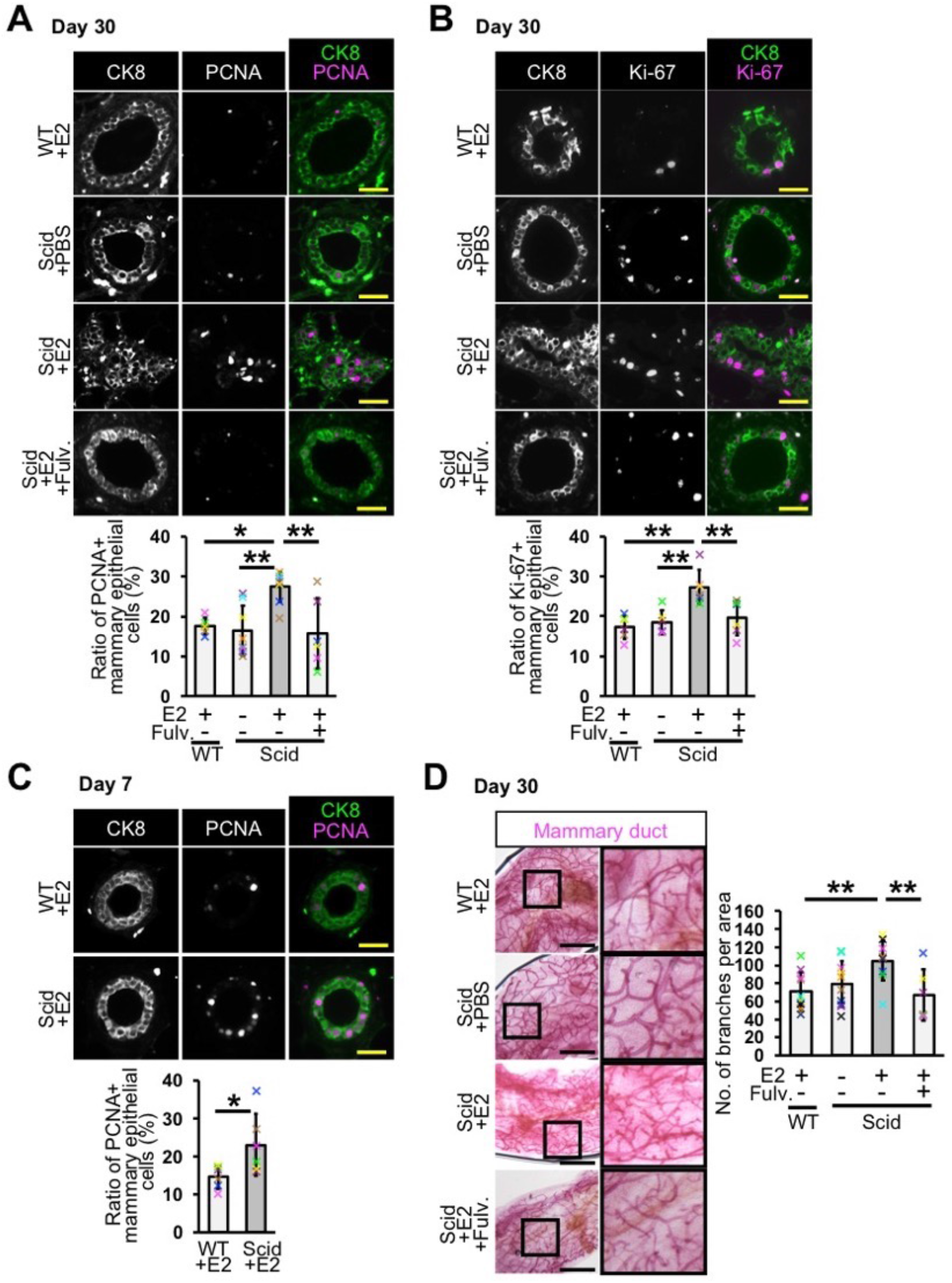
Estrogen administration promotes mammary epithelial cell proliferation in scid mice. A, PCNA positive mammary epithelial cells were detected at day 30. Ratios of the positive cells were analyzed (*n*= 6 mice in WT+E2 and Scid+E2+Fulv. groups, and *n*=8 mice in Scid+PBS and Scid+E2 groups, Tukey’s test). B, Ki-67 positive mammary epithelial cells were detected. Ratios of the positive cells were analyzed (*n*= 6 mice in WT+E2, Scid+PBS, Scid+E2 and Scid+E2+Fulv. groups, Tukey’s test). C, PCNA positive cells were analyzed in day 7-mice (*n*=6 mice, student’s *t*-test). D, Carmine Alum-staining was performed in mammary ducts of day 30-mice. Numbers of branches in 9 mm^2^ area close to lymph node were counted (*n*=10 mice in WT+E2, Scid+PBS and Scid+E2 groups, *n*=6 mice in Scid+E2+Fulv. group, Tukey’s test). Fulv.: fulvestrant. Scale bars indicate 30 μm (A-C) and 2 mm (D). *: *P*<0.05, **:*P*<0.01. Error bars represent standard deviation.

### Isoflavones prevent E2-induced dysplasia formation

Our dysplasia model system can be utilized to study breast cancer prevention. Isoflavones are flavonoids and are rich in soybean. Epidemiological studies have suggested that isoflavones have a protective effect against breast cancer (33). Genistein, an isoflavone, inhibited breast cancer cell growth (34). On the other hand, a study using breast cancer cells showed that genistein promoted tumor growth (35). Therefore, the effect of isoflavones in breast cancer is still controversial. To address this issue, we investigated the effect of isoflavones in our dysplasia model system. We used two isoflavones, (S)-equol and genistein. When E2-induced DNA double-strand breaks were analyzed, both isoflavones reduced E2-induced DNA damage (Fig. 4A), suggesting that isoflavones have a potential to inhibit E2 function.

**Fig. 4.**
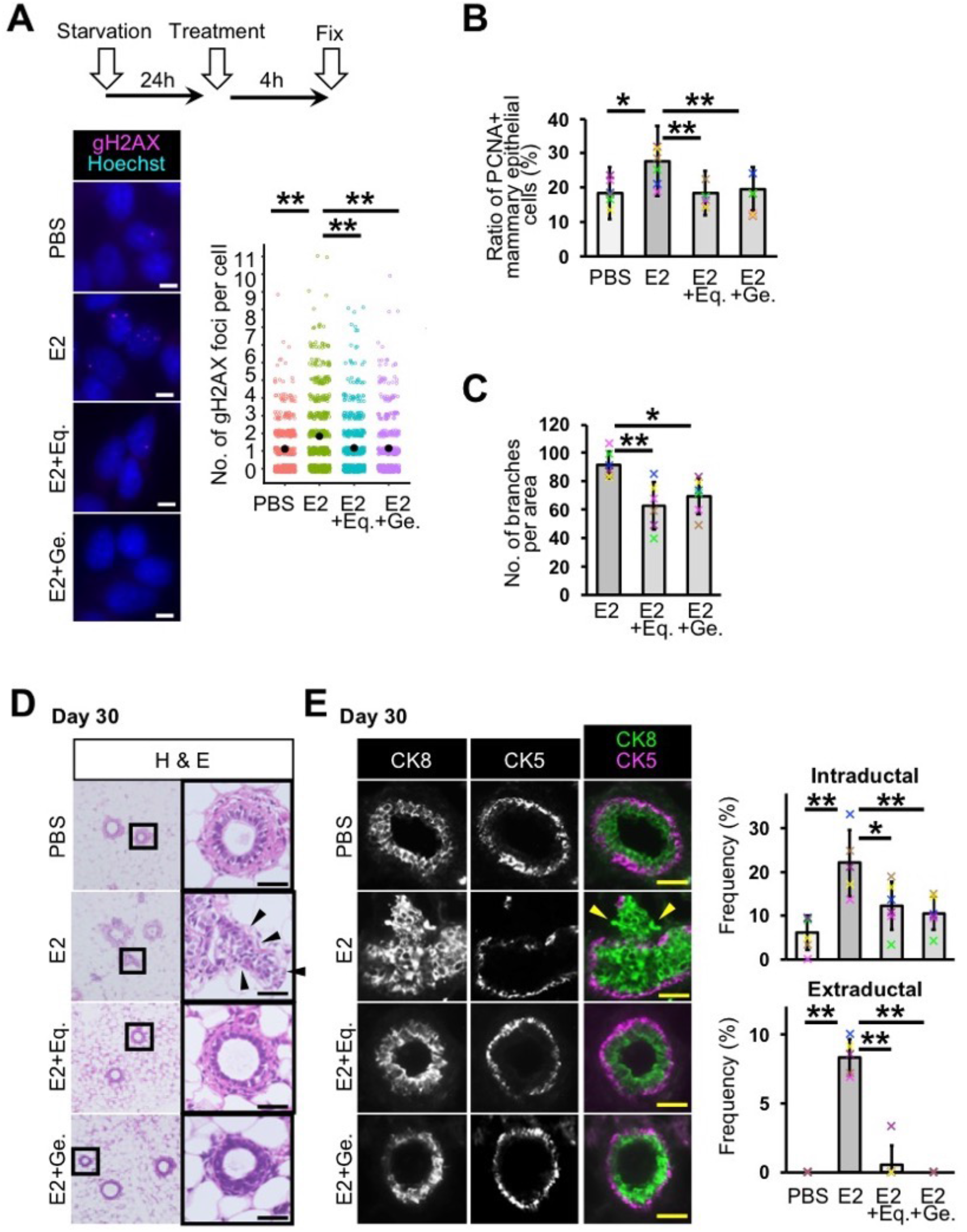
Isoflavones inhibit dysplasia formation by preventing the function of estrogen. A. DNA double-strand breaks were detected by gH2AX immunostaining. Numbers of gH2AX foci per cell were analyzed (jitter plot) (*n*=3 independent experiments, total 550~670 cells in each group, Mann-Whitney U test). B, Ratios of PCNA positive cells were analyzed (*n*=6 mice, Tukey’s test). C, Number of branches were analyzed in Carmine Alum-stained mammary glands (*n*=6 mice, Tukey’s test). D, Typical images of H&E staining are shown. E, Fluorescent images of CK8 and CK5 staining are shown. Mammary ducts with intraductal and extraductal expansions were quantified (*n*=6 mice in scid+PBS, scid+E2+Eq. and scid+E2+Ge. groups, *n*=5 mice in scid+E2, Tukey’s test and Mann-Whitney U test). Eq.: (S)-equol, Ge.: genistein. Scale bars indicate 10 μm (A) and 30 μm (E,F). *: *P*<0.05, **:*P*<0.01. Error bars represent standard deviation. Arrowheads indicate mammary epithelial cells in extraductal region (E,F).

The coadministration of E2 and isoflavones reduced cell proliferation of mammary epithelial cells *in vivo* (Fig. 4B, S4A). The number of branches was reduced by isoflavones (Fig. 4C,S4B). These results suggest that isoflavones were effective at preventing E2-induced cell proliferation. H&E staining showed that most of the mammary ducts of mice coadministered E2 and isoflavones had normal ductal structures and retained the biphasic mammary epithelial and myoepithelial pattern (Fig. 4D). The results of CK5 and CK8 immunostaining showed that the ratio of intraductal and extraductal expansion was reduced by isoflavone administration (Fig. 4E). These results indicate that isoflavones have the potential to prevent E2-induced dysplasia formation.

## Discussion

We demonstrated E2-induced mammary ductal dysplasia in mice, providing direct evidence that estrogen causes mammary ductal dysplasia. This study showed that reduced DNA repair capacity enhances estrogen signaling, which promotes dysplasia formation. In human, long-term estrogen exposure has the potential to cause breast cancer (17). On the other hand, DNA repair capacity is not maintained throughout life (36, 37). In the breast, human mammary epithelial cells from aged donors were more sensitive to mammography-induced DNA damage than the cells from young donors (38). Mammary epithelial cells from older women showed a delay of DNA double strand break repair, compared to those from younger women (39). Mutations in genes related to the DNA damage response/repair lead to the risk of developing breast cancer (40, 41). Our dysplasia-inducing model (i.e., 30-day E2 injection in mice with reduced DNA repair capacity) may mimic some of the situations of human breast tissue that are susceptible to mammary ductal dysplasia.

Although our dysplasia model uses an excess amount of E2 and mice with reduced DNA repair capacity, it will be useful to investigate the mechanism of dysplasia formation and to develop a method for dysplasia prevention. We showed dysplasia formation in wild-type strains by coadministration of the DNA-PK inhibitor and E2. These results indicate that our dysplasia model system can be utilized in various strains and genetically engineered mouse models, such as *Tp53*-knockout mice. For the breast cancer prevention study, we showed the first experimental evidence that isoflavones inhibited E2-induced dysplasia formation. In future, this dysplasia induction model system may contribute to the understanding of breast cancer tumorigenesis and to the development of breast cancer prevention.

## Methods

### Cell culture

MCF-7 cells were obtained from the American Type Culture Collection (Manassas, VA, USA). Short tandem repeat analysis was performed for cell authentication in July 2017 and the result showed no contamination and no alteration. Mycoplasma contamination was checked every 3 months by staining with Hoechst 33342 (Dojindo, 346-07951, Kamimashiki, Japan, 1/500 dilution) and no contamination was observed. Cells were maintained in Roswell Park Memorial Institute 1640 (RPMI-1640) medium containing 10% heat-inactivated fetal bovine serum (FBS), 100 units/mL Penicillin, 100 μg/mL Streptomycin and 1nM β-estradiol (E2) (Sigma, E2758, St. Louis, MO, USA) at 37°C and with 5% CO_2_. For drug selection to obtain infectants, 1 μg/mL puromycin was used. To obtain G1 phase cells For gH2AX staining experiments, cells were starved with phenol-red-free FBS-free RPMI-1640 medium containing 100 units/mL Penicillin and 100 μg/mL Streptomycin for 24 h. E2, progesterone (PG) (Sigma, P8783), (S)-equol (Cayman Chemical, 10010173, Ann Arbor, MI, USA) and genistein (Nagara Science, NH010302, Gifu, Japan) were solved in ethanol, diluted with phosphate buffered saline (PBS) and added to medium (final concentration: 10 nM). Fulvestrant (Sigma, I4409) was solved with ethanol, diluted with PBS and added to medium (final concentration: 100 nM). NU-7441 (AdooQ Bioscience, A11098, Irvine, CA, USA) was solved in DMSO, diluted with PBS and added to medium (final concentration: 0.5 μM). For mRNA quantification of ERα downstream genes, cells were cultured with phenol-red-free RPMI-1640 medium containing 10% charcoal-stripped FBS, 100 units/mL Penicillin and 100 μg/mL Streptomycin for 48 h, subsequently treated with or without 10 nM E2 for 6 h.

### Loss of function study

For short hair-pin RNA (shRNA) expression, a lentiviral vector pLKO.1 (Addgene, 8453, Cambridge, MA, USA) was used. Double-strand DNA oligo with shRNA sequence was cloned into the region between *Age*I and *Eco*RI sites of the vector. The target sequences were 5’-CCAGTGAAAGTCTGAATCATT-3’ (shPRKDC #2), 5’-CCTGAAGTCTTTACAACATAT-3’ (shPRKDC #4), 5’-GCTGCTGGAAGACGAAAGTTA-3’ (shPGR #1) and 5’-CAATACAGCTTCGAGTCATTA-3’ (shPGR #2). Control shRNA were 5’-CCTAAGGTTAAGTCGCCCTCG-3’ (shScr, control).

Lentiviral vector was co-transfected with lentiviral envelope and packaging plasmids, pMDLg/pRRE (Addgene, 12251), pMD2.G (Addgene, 12259) and pRSV-Rev (Addgene, 12253) at a ratio of 2.5:1.0:0.6:0.5 to Lenti-X 293T cells (Takara, 632180, Kusatsu, Japan). A transfection reagent, FuGENE 6 (Promega, E2691, Madison, WI, USA) was used. Lenti-X 293T cells were maintained with Dulbecco’s modified Eagle medium containing 10% FBS, 100 units/mL Penicillin and 100 μg/mL Streptomycin. One day after transfection, medium was changed to the medium for MCF-7 cells and incubated for 24~30 h. Medium containing lentiviral particles was filtered (0.22 μm pore size), added to MCF-7 cell culture with 6 μg/mL polybrene, and incubated for 48 h. For drug selection, infectants were treated with puromycin for 4 days.

### Messenger RNA quantification

Cells were cultured in 6-well plate or 6 cm dish. Cells were rinsed with cold PBS and treated with 300 μL of Trizol reagent (Thermo Fisher Scientific, 15596018, Waltham, MA, USA). Sixty μL of chloroform was added, mixed and stand for 5 min. After centrifugation at 4°C, supernatant was collected and purified with PureLink RNA Micro Kit (Thermo Fisher Scientific, 12183016).

Five hundred ng of Total RNA was used for complementary DNA (cDNA) synthesis. SuperScript III reverse transcriptase (Thermo Fisher Scientific, 18080044) was used. Synthesized cDNA was diluted with sterilized MilliQ water (1/10 dilution) for real-time polymerase chain reaction (PCR).

Real-time PCR was performed with a reagent, FastStart Universal SYBR Green Master (Sigma, 04 913 850 001, St. Louis, MO, USA). Signals were detected by StepOnePlus real-time PCR system (Thermo Fisher Scientific, 4376600) with StepOne software ver2.2.2.

Primer sequences were: *EF1A1* (internal control) forward: 5’-AAATGACCCACCAATGGAAGCAGC-3’ reverse: 5’-TGAGCCGTGTGGCAATCCAATACA-3’, *PRKDC* forward: 5’-CGCCGTGTGAATATAAAGATTGG-3’ reverse: 5’-CGTGACTGTTTCAGTACGATTAG-3’, *GREB1* forward: 5’-CTGCTGTACCTCTGTGACTCTT-3’ reverse: 5’-GTCCTGACAGATGACACACAAC-3’, *TFF1* forward: 5’-CCCTGGTCCTGGTGTCCAT-3’ reverse: 5’-AGCAGCCCTTATTTGCACACT-3’, *PGR* forward: 5’-CACAGCGTTTCTATCAACTTACAA-3’ reverse: 5’-CCGGGACTGGATAAATGTATTC-3’.

### Immunostaining in cell culture

Cells were plated onto an 8-well chamber slide (Matsunami glass, SCS-N08, Kishiwada, Japan) with 400 μL medium, and cultured for 2 days. For fixation, a half of the medium was removed, and 200 μL of 4% paraformaldehyde (PFA) in PBS was added (final 2% PFA). After 10~15 min fixation at room-temperature, cells were washed with PBS containing 0.05% Tween-20 (PBS-T). Permeabilization was performed with PBS containing 0.1% Triton-X 100 for 15 min at room-temperature. Cells were washed with PBS-T twice. Blocking was performed with 5% goat serum-containing PBS-T for 1 h at room-temperature. Cells were incubated with primary antibody in blocking solution for overnight (15~20 h) at 4°C. Primary antibody was anti-gH2AX antibody (S139) (Cell Signaling Technology, 2577S, Danvers, MA, USA, 1/200 dilution). Cells were washed with PBS-T 3 times and incubated with secondary antibody in blocking solution for 1~2 h at room-temperature. Secondary antibody was goat anti-rabbit IgG antibody conjugated to Alexa Fluor 546 (Life Technologies, A11010, 1/1000 dilution). Cells were washed with PBS twice and counterstained with Hoechst 33342 for 30 min at room-temperature. Cells were washed with PBS, dried and mounted with Fluoromount-G (SouthernBiotech, 0100-01, Birmingham, AL, USA).

### Mouse experiments

Female C.B17/Icr wildtype (C.B17/Icr-scidJcl +/+) and scid (C.B17/Icr-scidJcl scid/scid) mice were purchased from CLEA Japan (Tokyo, Japan) (6~8-week-old). Six-week-old female C57BL/6J mice were purchased from Japan SLC (Hamamatsu, Japan). Intraperitoneal injection was performed with 30G needle in the morning. Injected reagents were E2 (6 μg/day), PG (6 μg/day), Fulvestrant (100 μg/day), NU-7441 (100 μg/day), (S)-equol (6 μg/day) and genistein (6 μg/day). Mice were euthanized by cervical dislocation, and mammary glands were isolated. For 30-day samples, mammary glands were isolated at 24 h after final injection. For measurement of E2 serum concentration after administration, 5-week-old female mice were ovariectomized to eliminate endogenous E2. After 5 weeks, E2 was injected intraperitoneally, and blood samples were collected from tail vein. E2 concentration was measured by using E2 ELISA(EIA) kit (Calbiotech, ES180S-100, El Cajon, CA, USA). The animal experiments were approved by the Animal Research Committee of Kyoto University, number MedKyo17554 and MedKyo18321. All animals were maintained according to the Guide for the Care and Use of Laboratory Animals (National Institute of Health Publication).

### Immunostaining in mammary gland

For cryo-section, isolated mammary gland was fixed with 4% PFA in PBS shortly (4 °C, 15 min, rocking). Sample was washed with PBS 3 times and incubated with 30% sucrose in PBS for 1~2 h at room-temperature. Samples was embedded in OCT compound (Sakura Finetek, 4583, Tokyo, Japan), and frozen with liquid nitrogen. Ten μm cryo-section was cut at −50°C, and dried. Dried section was fixed with 4% PFA for 3 min at room-temperature and rinsed with PBS. For paraffin-section, mammary gland was fixed with 10% formaldehyde neutral buffer solution for longer than 24 h at room-temperature, dehydrated and embedded in paraffin. Three μm paraffin-section was de-paraffinized, washed with PBS and rinsed with H_2_O. For heat-induced epitope retrieval, specimen was put into boiling sodium citrate buffer (10 mM sodium citrate, 0.05 % Tween-20, pH 6.0), incubated for 40 min and cooled for 20 min.

Specimen was washed with PBS-T 3 times and blocked with blocking solution (5% goat serum in PBS-T) for 1 h at room-temperature. Specimen was incubated with primary antibody in blocking solution for overnight (15~20 h) at 4 °C. Primary antibodies were anti-gH2AX antibody (Cell Signaling Technology, 2577S, 1/200 dilution), anti-CK8 antibody (Developmental Studies Hybridoma Bank, TROMA-I, Iowa City, IA, USA 1/200 dilution), anti-CK5 antibody (Abcam, ab75869, 1/200 dilution), anti-PCNA antibody clone PC10 conjugated to Alexa Fluor 647 (BioLegend, 307912, San Diego, CA, 1/20 dilution), anti-Ki-67 antibody D3B5 (Cell Signaling Technology, 12202S, 1/400 dilution) and anti-ERα antibody (Millipore, 06-935, 1/400 dilution) and anti-ERα antibody SP1 (NeoMarkers, RM-9101-S0, Fremont, CA, USA, 1/100 dilution). Specimen was washed 3 times with PBS-T and incubated with secondary antibody in blocking solution. Secondary antibodies were biotinylated goat anti-rabbit IgG (Vector, BA-1000, Burlingame, CA, USA, 1/200 dilution), goat anti-rat IgG conjugated to Alexa Fluor 488 (Cell Signaling Technology, 4416S, 1/1000 dilution) and goat anti-rabbit IgG antibody conjugated to Alexa Fluor 546 (Life Technologies, A11010, 1/1000 dilution).

For DAB colorimetric detection, after secondary antibody reaction, specimen was washed with PBS 3 times and incubated with ABC kit (Vector, PK-6101). Specimen was washed with PBS and rinsed with H_2_O. Specimen was washed with H_2_O and counterstained with hematoxylin. Specimen was rinsed with H_2_O and washed with H_2_O for 10 min. Dehydrated specimen was mounted with Malinol (Muto Pure Chemicals, 2009-1, Tokyo, Japan).

For immunofluorescence, after secondary antibody reaction, specimen was washed with PBS twice and counterstained with Hoechst 33342 for 30 min at room-temperature. Specimen was washed with PBS, dried and mounted with Fluoromount-G.

### Carmine Alum-staining

Isolated mammary gland was foxed with 4% PFA for 2 h at 4°C, washed with PBS twice and washed with H_2_O. Sample was incubated with Carmine Alum staining solution (2 mg/mL carmine, 5 mg/mL aluminum potassium sulfate, a small amount of thymol) for overnight (20~24 h) at room-temperature. After staining, sample was washed with 70% ethanol for 1 h at room-temperature, 95% ethanol for 1 h at room-temperature and 100% ethanol for 1 h at room-temperature. Subsequently sample was cleared with xylene overnight (16~20 h) at room-temperature. Xylene was replaced to methyl salicylate for storage.

### microscopy

Images of H&E staining, immunostaining and *in situ* hybridization were collected at room temperature with an all-in-one microscope BZ-9000 (Keyence, Osaka, Japan) equipped with a 20x plan apochromatic objective lens (NA: 0.75), a x40 plan apochromatic lens (NA: 0.95) and a x100 plan apochromatic lens (NA: 1.40), and BZ-II Viewer software (Keyence). Hoechst 33342 signal was excited by 340-380 nm light and detected with 435-485 nm light. Alexa Fluor 488 signal was excited by 450-490 nm light and detected with 510-560 nm light. Alexa Fluor 546 signal was excited by 527.5-552.5 nm light and detected with 577.5-632.5 nm light. Alexa Fluor 647 signal was excited by 590-650 nm light and detected with 662.5-737.5 nm light.

Images of carmine alum-stained mammary gland were collected at room-temperature with a stereoscope, SMZ800 (Nikon, Tokyo, Japan) and Digital Sight DS-Fi1 (Nikon)

### Statistical analyses

Numbers of gH2AX foci of cultured cells were counted manually and analyzed by Mann-Whitney U test. Delta Ct values of real-time PCR experiments were normalized by the mean values of the control groups and analyzed by student’s *t*-test and Tukey’s test. Numbers of mammary epithelial cells having more than 5 gH2AX foci were counted manually and analyzed by Tukey’s test. Values of E2 serum concentration were normalized by the mean values of 0 h samples and analyze by student’s *t*-test between wildtype and scid mice in each time point. Numbers of Mammary ducts with intraductal and extraductal expansions were analyzed by Tukey’s test and Mann-Whitney U test. Numbers of mammary epithelial cells having immunostaining signals of PCNA, Ki-67 and ERα were counted manually, and analyzed by Tukey’s test and student’s *t*-test. Number of branches were analyzed by Tukey’s test and student’s *t*-test. *P*<0.05 was considered to be statistically significant.

## Acknowledgement

We thank Dr. Makoto Noda, Dr. Yo-ichi Nabeshima, and Mr. Akihiro Nakamura for fruitful discussion. We thank Dr. Shin-ichiro Imai for writing assistance. We thank Dr. Yasuhiko Kawakami for sharing the information about experiments. We thank Ms. Sunao Tanaka, Dr. Noriko Senda and Ms. Remi Akagawa for technical assistance. We thank Medical Research Support Center, and Center for Anatomical, Pathological and Forensic Medical Researches, Graduate School of Medicine, Kyoto University for technical assistance. Short tandem repeat analysis for cell authentication was performed by BEX Co., Ltd. The manuscript was proofread by *American Journal Experts*. We thank the Developmental Studies Hybridoma Bank developed under the auspices of the NICHD and maintained by the University of Iowa. Financial supports were provided by Taiho Pharmaceutical Co., Ltd., Cactus Communications K.K., Sanwa Shurui Co. Ltd. and JSPS KAKENHI Grant Number 19K07665.

## Supplemental figures

**Fig. S1.**
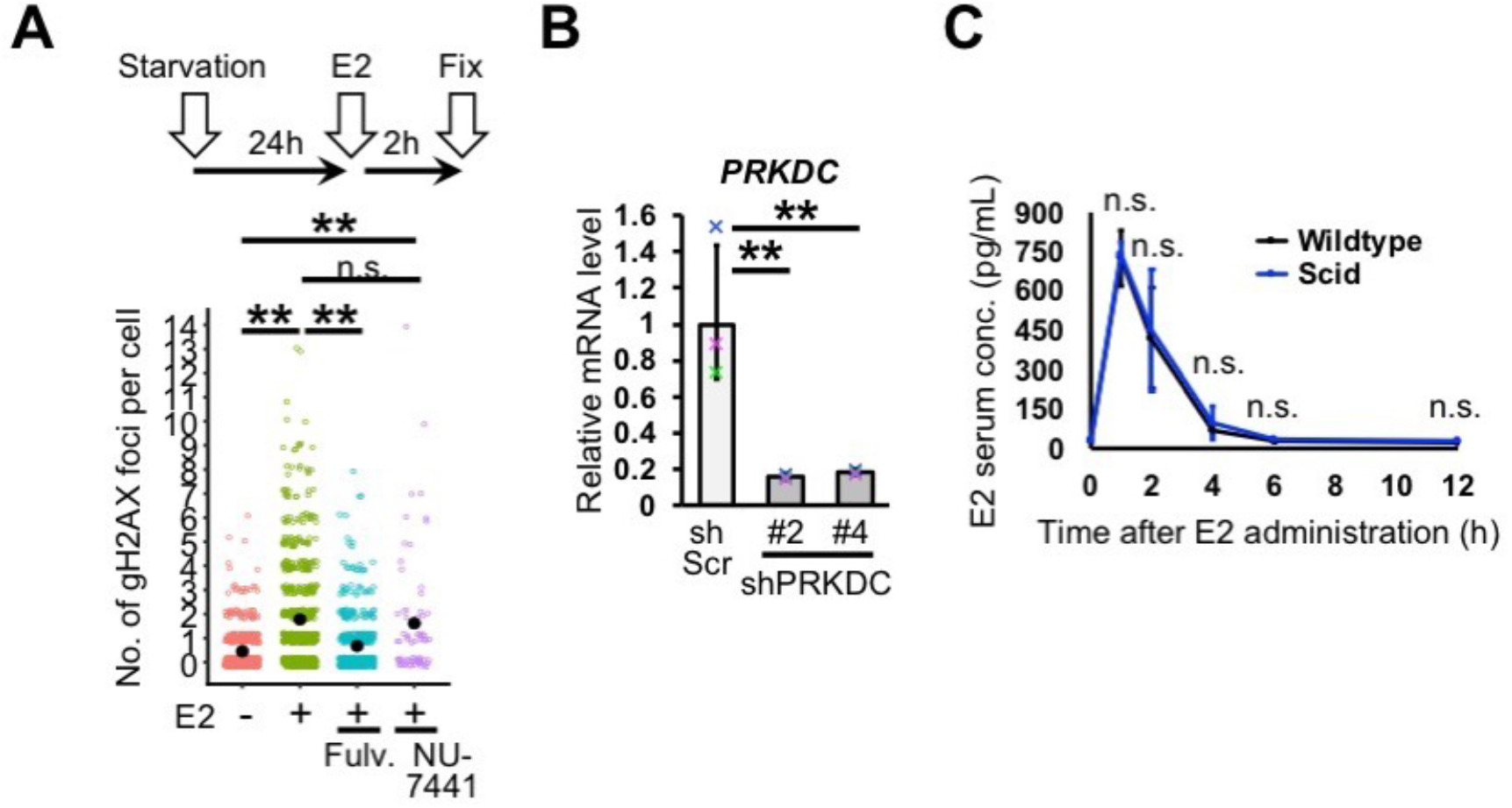
Estrogen administration induces DNA double-strand break. A, DNA double-strand breaks were detected in MCF-7 cells. Numbers of gH2AX foci per cell were graphed (jitter plot). Black dots indicate mean values (total 70~760 cells in each group, Mann-Whitney U test). Fulv.: fulvestrant, an estrogen receptor inhibitor. NU-7441, a DNA-PK inhibitor. B, *PRKDC* gene was knocked-down (*n*=3 experiments, student’s *t*-test to shScr control). MCF-7 cells were used. C, E2 serum concentration was measured (*n*=3 mice, student’s *t*-test in each time point). n.s.: not significant, **:*P*<0.01. Error bars represent standard deviation.

**Fig. S2.**
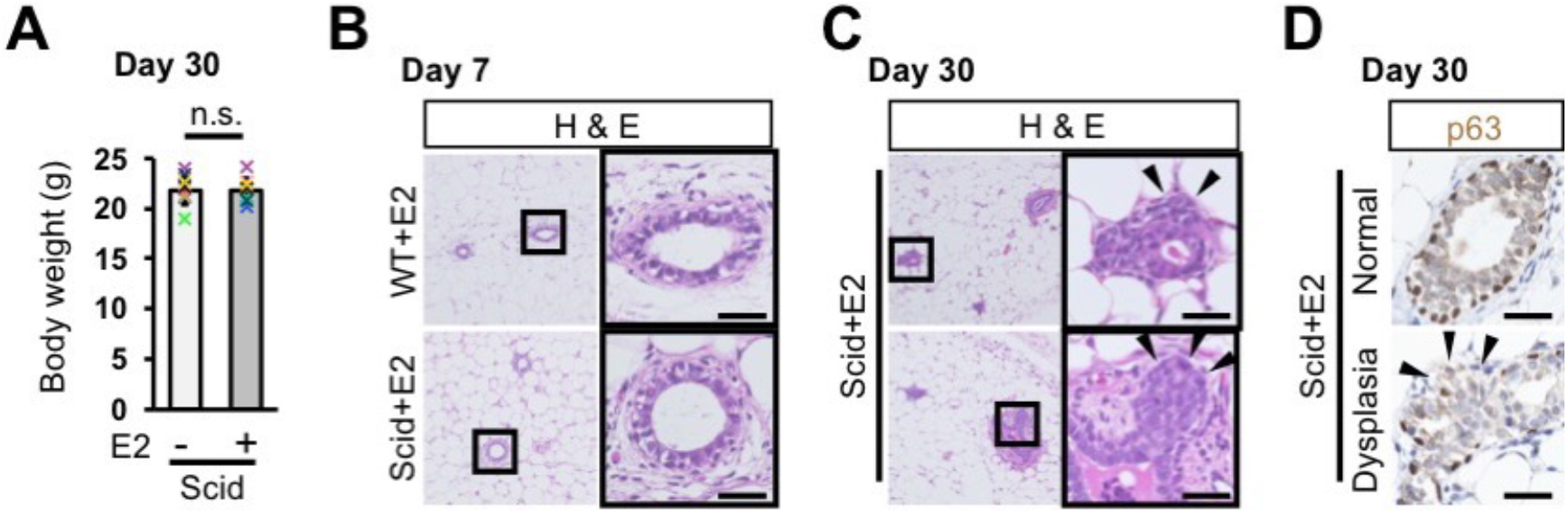
Long-term estrogen administration causes dysplasia. A, Body weight was measured at day 30 (*n*=10 mice, student’s *t*-test). B, Images of H&E staining of mammary glands are shown. Mammary glands were isolated at day 7. C, Additional to Fig. 1D, H&E images of mammary glands of E2-adinistered scid mice are shown. Arrowheads indicate mammary epithelial cells in extraductal region. D, The myoepithelial marker, p63, was stained. Arrowheads indicate a region lost myoepithelial cells. Scale bars indicate 30 μm. n.s.: not significant. Error bars represent standard deviation.

**Fig. S3.**
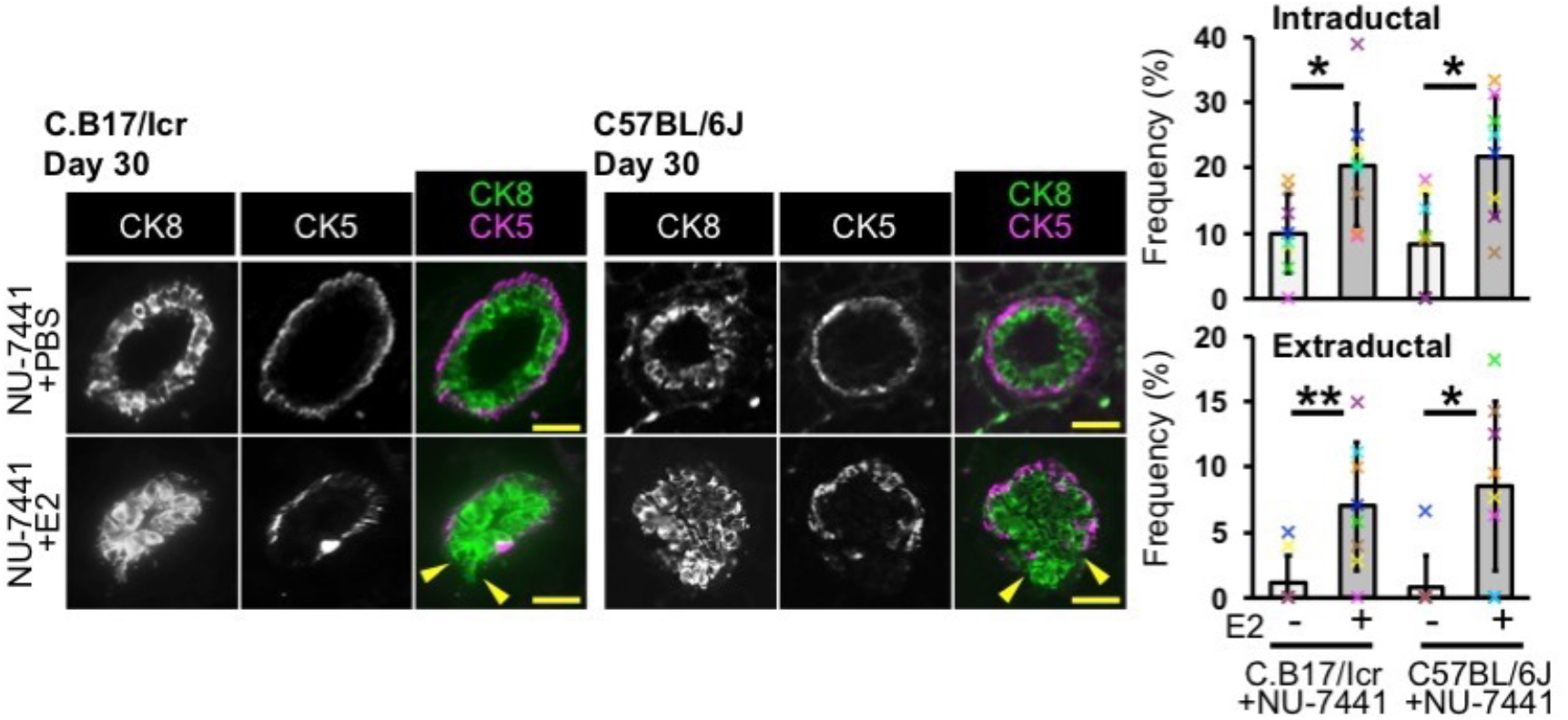
Dysplasia is induced by the combination of E2 administration and DNA-PK pharmacological inhibition. Fluorescent images of CK8 and CK5 staining are shown. A DNA-PK inhibitor, NU-7441, was administered to wildtype stains, C.B-17/Icr and C57BL/6J, in combination with or without E2. Mammary ducts with intraductal and extraductal expansion were quantified (*n*=8 mice, Mann-Whitney U test in each strain). Arrowheads indicate mammary epithelial cells in extraductal region. Scale bars indicate 30 μm. *: *P*<0.05, **:*P*<0.01. Error bars represent standard deviation.

**Fig. S4.**
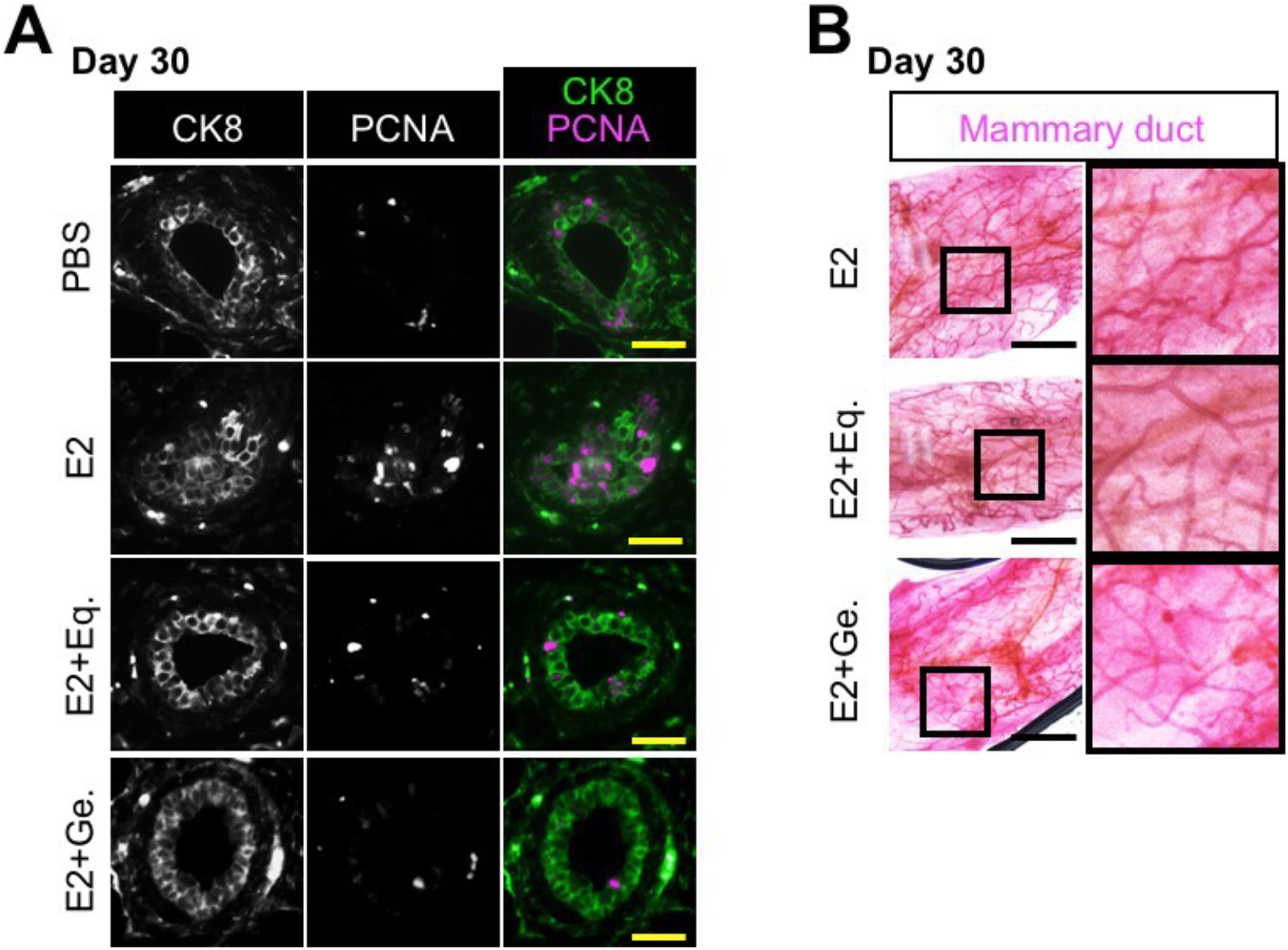
Isoflavones inhibits estrogen-promoted cell proliferation in mammary gland. A, Typical images of PCNA staining are shown. Quantification is shown in Fig. 4C. B, Typical images of Carmine Alum-staining are shown. Numbers of branching are graphed in Fig. 4D. Eq.: (S)-equol, Ge.: genistein. Scale bars indicate 30 μm (A) and 2 mm (B).

